# Growth but not corticosterone, oxidative stress or telomere length is negatively affected by microplastic exposure in a filter-feeding amphibian

**DOI:** 10.1101/2025.01.14.632987

**Authors:** Colette Martin, Katharina Ruthsatz, Ivan Gomez-Mestre, Pablo Burraco

**Affiliations:** Zoological Institute, Technische Universität Braunschweig, Mendelssohnstraße 4, 38106 Braunschweig, Germany; Institute of Biodiversity, Animal Health and Comparative Medicine, University of Glasgow, Glasgow, UK; Ecology, Evolution, and Development Group, Department Ecology and Evolution, Doñana Biological Station, CSIC, 41092 Seville, Spain

**Keywords:** *Xenopus laevis*, plastic pollution, amphibian decline, ageing, redox status, glucocorticoids

## Abstract

Microplastics (MPs) are of increasing global concern for species inhabiting aquatic habitats. However, the mechanisms behind animal responses to MPs need comprehensive exploration. Amphibians are the most threatened vertebrate group with most species having a complex life cycle, commonly with an aquatic larval stage. Here, we investigated whether exposure to an environmentally-relevant concentration of MPs affects the growth of filter-feeding larvae of the African clawed frog (*Xenopus laevis*), and the consequences for their stress physiology (corticosterone (CORT) levels), or health and ageing physiology (oxidative stress and telomere length). We conducted a 3×2 experiment with three levels of fibre exposure (fibres absent -control-, and MP and cellulose fibre treatments), and two stress levels (CORT absent –control-, and CORT present simulating a stressful condition). We observed a negative impact of MP exposure on larval growth; however, this did not alter the CORT levels, oxidative stress or telomere length. Our study shows that realistic concentrations of MPs is not enough to induce major alterations on the stress or health and ageing physiology of a filter-feeding amphibian. Whether compensatory growth responses during the post-metamorphic stages could lead to detrimental effects later in life should be explored in amphibians and other organisms with complex-life cycles.

## 1. Introduction

Human-driven activities expose wildlife to a myriad of stressors, including altered temperatures, habitat loss, and pollution (Lee et al., 2023). These conditions have the potential to impair vital processes such as metabolism, growth, development, or reproduction, which finally can compromise organismal health and survival (Román-Palacios & Wiens, 2020; Sheridan & Bickford, 2011). Consequently, anthropogenic disturbances may adversely affect fitness and lead to population declines, thereby contributing to the current global biodiversity loss (Díaz et al., 2019). Some physiological pathways have been recently presented as markers of individual fitness and health or ageing (e.g. Angelier & Wingfield, 2012; Burraco et al., 2020; Lemaître et al., 2022; Schoenle et al., 2021), and thus they can improve our understanding of eco-toxicological processes and advance conservation management plans.

A central physiological mechanism mediating animal responses to habitat changes is the hypothalamic–pituitary–adrenal/interrenal (HPA/I) axis (Crespi et al., 2013; Sapolsky et al., 2000). When internal homeostasis is disrupted, the activation of this neuroendocrine stress axis ultimately results in the secretion of glucocorticoids (GCs). Cortisol is the predominant GC in most primates and teleost fish, and corticosterone (CORT) in most amphibians and sauropsids (Angelier & Chastel, 2009; Glennemeier & Denver, 2002; Schreck & Tort, 2016; Suarez-Bregua et al., 2018). The release of GC hormones mobilises energy stores to meet increased metabolic demands in order to regulate metabolism and nutrient homeostasis (Kirschman et al., 2017; Sapolsky et al., 2000). In addition to energy-related functions, GCs play a pivotal role in developmental processes across many taxa (rev. in Crespi et al., 2013). For instance, in animals with at least two discrete life stages, GCs can plastically accelerate development and growth, allowing the transition to the next life stage to escape suboptimal habitats (Crespi & Warne, 2013; Denver, 2009; Gomez-Mestre et al., 2013). GC release and subsequent enhanced development can include cascading effects at different molecular levels such as the induction of an oxidative state (Costantini, 2014, 2019), which has the potential to damage the structure and function of molecules within the cell, including DNA (Halliwell, 2007; Monaghan et al., 2009). Particularly, oxidative stress is known to lead to telomere shortening (Armstrong & Boonekamp, 2023; Chatelain et al., 2020). When telomeres become critically short, cell apoptosis is induced (Lin & Epel, 2022). Therefore, telomere shortening is considered a hallmark of cellular and organismal ageing (López-Otín et al., 2023) and can predict fitness and survival odds (Eastwood et al., 2023; Q. Wang et al., 2018; Wilbourn et al., 2018). Integrating these health and ageing-related parameters into ecotoxicology and evolution will therefore enable us to better understand the molecular processes governing human-driven life history shifts.

Environmental pollution resulting from, for instance, industrial waste, agricultural chemicals or urban runoff, is dramatically pushing many natural populations to the brink of extinction (Noyes & Lema, 2015; Sigmund et al., 2023; Wake & Vredenburg, 2008). Aquatic habitats are particularly threatened by pollutants since these environments act as a sink for almost all kind of contaminants (Kumar et al., 2021; Mushtaq et al., 2020; Rzymski et al., 2017). Among several other pollutants, the presence of microplastics (MPs) is an increasing global concern for aquatic organisms due to large production volumes, continuous and worldwide release, and long-term environmental persistence in ecosystems (Li et al., 2018; PlasticsEurope, 2022; Windsor et al., 2019). MPs are defined as synthetic polymer particles with diameters ranging from 1 μm to 5 mm (Hartmann et al., 2019; Thompson et al., 2004). The source of MPs is diverse as they derive from either primary plastics used for purposes such as personal care products, plastic production pellets, and textiles, or from secondary plastics such as debris of plastic items, fishing nets, or tires (Horton et al., 2017). Despite being most abundant in the marine environment (rev. in Wang et al., 2022), freshwater environments have been identified as a key pathway in the transport of MPs from terrestrial to marine ecosystems (rev. in Stanton et al., 2020). Given their size range, MPs fall within the prey range for a variety of aquatic animals and thus they can be mistakenly ingested (Franzellitti et al., 2019) possibly resulting in adverse organismal effects, including reductions in growth, reproduction, and survival (rev. in Li et al., 2023; Prokić et al., 2019). However, the mechanisms driving these major effects of MPs are not very well understood in many taxa.

Amphibian larvae represent an ideal study system to investigate the mechanistic consequences of MP pollution on aquatic organisms due to their non-selective feeding mode, hormone-regulated development, semi-permeable skin and limited dispersal capacity (Gonçalves et al., 2024; Ruthsatz & Glos, 2024; Shi, 2000). Amphibians coping with stressors activate the HPI axis which leads to CORT and thyroid hormones release (Kirschman et al., 2017; Kulkarni & Buchholz, 2014). At the larval stage, higher CORT levels are known to accelerate development and growth rates, then allowing the transition to the terrestrial post-metamorphic stage to avoid suboptimal habitat conditions (Crespi & Denver, 2005; Crespi & Warne, 2013; Denver, 2009). Amphibian larvae of many species are known to express developmental plasticity in response to conditions such as pond drying (e.g. Kulkarni et al., 2017; O’Regan et al., 2014; Székely et al., 2017), increased water temperature (Sinai et al., 2022) or pathogen presence (Warne et al., 2011), all responses resulting in higher CORT levels. Also, altered developmental trajectories can include trade-offs since organisms developing faster often experience reductions in body condition, locomotor performance, or immunocompetence (Burraco et al., 2022b; Gervasi & Foufopoulos, 2008; Ruthsatz et al., 2019a; Ruthsatz et al., 2020), as well as timing of sexual maturation or even survival odds (Burraco et al., 2023b). These processes can contribute to the fact that approximately 41% of all amphibian species are threatened with extinction, with climate change, habitat loss and environmental pollution acting as major drivers of those declines (Luedtke et al., 2023; Stuart et al., 2004).

In relation to MP contamination, research has shown that polymer ingestion has detrimental effects on amphibian larval growth (Balestrieri et al., 2022; but not: De Felice et al., 2018), behaviour (da Costa Araújo & Malafaia, 2020; but not: Scribano et al., 2023), body condition (Boyero et al., 2020), developmental rate (Ruthsatz et al., 2022, 2023),and metabolism (Ruthsatz et al., 2023b). Furthermore, MP accumulation has been found in the digestive tract of amphibians following ingestion (Hu et al., 2016), which may result in carry-over effects, particularly in filter-feeding species potentially ingesting large amounts of MPs present in the water body. However, only three recent studies have addressed the impact of MPs on some physiological markers of health and stress in amphibians (da Costa Araújo et al., 2023; Attademo et al., 2023; Ruthsatz et al., 2023b), and thus further comprehensive approaches will finally determine the actual impact of MPs on the condition of amphibians and other aquatic organisms.

Here, we investigated whether MP ingestion has consequences for CORT levels, oxidative stress, and telomere length in larvae of the filter-feeding African clawed frog (*Xenopus laevis*). The experimental set-up consisted of a 3×2 design, including three levels of fibre exposure (a no fibres control, and MP and cellulose fibre treatments), and two levels of simulated stress (presence/absence of exogenous CORT). We predicted that MP ingestion would decrease larval growth and that it would result in impaired physiology, including increased CORT levels and oxidative stress (or, alternatively, buffered oxidative stress through increased antioxidant responses), and telomere shortening.

## 2. Materials and Methods

### 2.1 Experimental design and animal husbandry

Two egg clutches of *Xenopus laevis* were obtained from the Andalusian Center for Development Biology (CABD, Pablo de Olavide University and CSIC), and kept at 24°C in a single bucket filled with de-chlorinated water. When larvae reached the developmental stage NF40 (free-swimming stage; Nieuwkoop P., 1994), we randomly allocated one individual to each of 102 round containers filled with 2.6 L of de-chlorinated water (i.e., 17 tadpoles per treatment group). The experiment was conducted in a climate chamber set to 24 °C and with a 14:10 h light:dark photoperiod. Larvae were fed a protein rich powdered fish food consisting of 50:50 sera micron powder and spirulina (Sera, Germany). This food is confirmed to be free of MPs and falls within the size range associated with MPs and cellulose (Ruthsatz et al., 2023b). Food was provided *ad libitum*, and sufficient rations were provided twice daily to ensure a constant and abundant food supply throughout the experiment. Larvae were maintained under these control conditions until they reached NF46 (compact intestine formed; Nieuwkoop P., 1994), the point at which we started exposing larvae to the different treatments.

### 2.2 Fibre and CORT exposure

We used polyethylene microplastic (MP) 34–50 μm particles (Sigma-Aldrich; polyethylene powder, CAS number 9002-88-4). Polyethylene is one of the most commonly used polymers for creating plastic materials, also known to be a significant source of MPs in the wild (Horton et al., 2017; Karaoğlu & Gül, 2020). Amphibians, particularly tadpoles, are exposed to polyethylene MPs worldwide (Hu et al., 2018; Karaoğlu & Gül, 2020). Previous research has investigated the effect of polyethylene on amphibian life-history and health (da Costa Araújo & Malafaia, 2020; Ruthsatz et al., 2022; Ruthsatz et al., 2023b). We chose a MP concentration of 60 mg/L, following the procedure adapted from da Costa Araújo et al. (2020a). This concentration, as reported by Ruthsatz et al. (2023), corresponds to a particle density of 1.0356–1.0675 × 10^7^ particles per litre. This density falls within the environmentally relevant range of surface water contamination with MP (Koelmans et al., 2019), and serves as an indicator of high pollution levels (da Costa Araújo et al., 2020a; da Costa Araújo et al., 2020b). Cellulose (Sigma-Aldrich; cellulose powder, CAS number 9004-34-6, particle size: 51 μm) was used at a concentration of 60 mg/L as a natural fibre control treatment in this experiment, *i.e.,* to control for a possible effect of exposure to fibres within the same size range (∼50 μm in our case), but not of a plastic nature (Buss et al., 2021).

When tadpoles reached developmental stage NF46, the MP and cellulose fibres were added directly to the water in the experimental containers (i.e., 60 mg/L x 2.6 L water = 156 mg fibres per container). Air stone bubble diffusers ensured both constant and effective water aeration and the continuous dispersion of fibres within the water, preventing particle settling and the formation of a MP film on the water surface (Ruthsatz et al., 2022; Ruthsatz et al., 2023b). Five days after the start of fibre exposure, we added 100 nM of CORT to half of the buckets (N=51). The water was fully renewed every other day to ensure the desired CORT concentration. During each water change, we used a separate net and bucket for each treatment, we wore cotton material clothing (i.e., polymer-free) and bright blue nitrile gloves, and hair was tied back, to prevent cross-contamination. An air purifying system (Philips AC2889/10, CADR 333 m^3^ × h^−1^) was used at all times in the experimental climate chamber to filter possible air contamination.

### 2.3 Sampling

Larvae were euthanised when they reached developmental stage NF57 (i.e. all five toes separated; Nieuwkoop et al., 1994) via immersion in a lethal solution of tricaine methanesulfonate (2 g/L MS-222, Ethyl 3-amino-benzoate methanesulfonate; Sigma-Aldrich), buffered with 200 mg/L sodium bicarbonate (Cecala et al., 2007). We blotted dry and weighed each individual to the nearest 0.0001g using a high precision balance (Ohaus VP-114CN Voyager Analytical Balance, Spain). We measured snout-vent length (SVL) and total length (TL) to the nearest 0.5 mm with a calliper. Then, we dissected the liver and gut, blotted dry, and weighed (only the gut) separately to the nearest 0.0001g using a high precision balance (Ohaus VP-114CN Voyager Analytical Balance, Spain). The collected livers and guts were placed separately in 1.5 mL Eppendorf tubes, snap frozen, and stored at −80 °C until used for telomere length quantification. The body remnants and tails were stored in 1.5 mL Eppendorf tubes and stored at −80 °C for subsequent oxidative stress and CORT analyses.

### 2.4 Determination of physiological parameters

#### 2.4.1 Corticosterone (CORT) assay

Hormone extraction took place in August 2023. We thawed and weighed each tail sample to the nearest 0.0001 g followed by homogenisation in 13×100 mm glass tubes with 500 µL PBS buffer (AppliChem Panreac, Germany) using a homogeniser at ∼17,000 rpm (Miccra D-1, Germany). In order to collect the sample residue, we washed the tissue blender into the tube with additional 500 µL PBS buffer. Then, the blender was cleaned with ddH20, and 96 % EtOH to avoid cross contamination. After homogenisation, we added 4 mL of a 30:70 petroleum ether:diethyl ether dissolvent mixture (both from Sigma-Aldrich, Germany) to each sample. Samples were vortexed for 60 s and subsequently centrifuged at 1,800 g and 4 °C for 15 min. Then, we snap-froze samples in a dry ice ethanol bath for 5 min, collected the resulting top organic layer containing CORT, and placed each sample in a new 13×100 mm glass tube. We repeated all steps after homogenisation to ensure maximum CORT extraction, and we pooled each recovered ether fraction into a single tube. We evaporated each sample with the help of a sample concentrator (Techne FSC400D; Barloworld Scientific, United Kingdom) consisting of a constant but gentle nitrogen flow. Finally, we resuspended lipids in 350 μL EIA buffer (Assay buffer, DetectX Corticosterone ELISA kit, K014-H5, Arbor Assays, Ann Arbor, MI, USA) with the help of a vortex, then tubes were sealed with parafilm and incubated overnight in a fridge at 4 °C.

We measured CORT levels using DetectX Corticosterone ELISA (Enzyme Immunoassay) kits from Arbor Assays (K014-H5, Ann Arbor, MI, USA). ELISA assays have been validated and successfully used for CORT detection in several amphibians (e.g., *Lithobates sylvaticus*, Gavel et al., 2019; *Rana arvalis*, Mausbach et al., 2022; *R. temporaria*, Burraco et al., 2017; Ruthsatz et al., 2023a; *X. laevis*, Ruthsatz et al., 2023b). We used the 100 µL assay format for standard preparations and assays. CORT concentration was measured in triplicates for all samples on 96-well plates according to the kit’s instructions. The plates were read with a multimode microplate reader (MB-580 3, Heales) at 450 nm. In total, we ran 4 plates. MyAssays online tools were used to calculate the hormonal concentration of samples based on calibration standards provided with the DetectX kit (https://www.myassays.com/arbor-assays-corticosterone-enzyme-immunoassay-kit-improved-sensitivity.assay). A new standard curve for calculation of the results was run for each plate. The mean coefficient of variation of triplicates for all samples was 19.91 %. Intraplate variation was overall 24.6 % and interplate variation was on average 30.01 %. We kept all the measurements of triplicates with a coefficient of variation lower than or equal to 30.0% or with an absolute difference between mean and median lower than 2.5 pg (Ruthsatz & Rico-Millan et al., 2023). Each plate also included a negative control with no CORT nor antibody. The average background CORT recorded in our negative control samples was subtracted from hormone samples. Average *R*^2^ for the 4PLC fitting curve was 0.994.

#### 2.4.2 Oxidative stress

We quantified the activity of four antioxidant enzymes (catalase, glutathione reductase, glutathione peroxidase, and superoxide dismutase), lipid peroxidation (malondialdehyde levels) and antioxidant status (reduced-to-oxidised glutathione ratio). The remnants of larvae (i.e., after collecting the tail for CORT quantification, and the gut and liver for telomere length measurements) were immersed in a buffered solution that inhibits proteolysis (Tris HCl 100nM pH 7.8, EDTA 0.1 mM and 0.1% Triton X-100) using a 1:4 (weight:volume) proportion, then were homogenised at 35,000 rpm with a Miccra homogeniser (Miccra D-1). We centrifuged the homogenates at 20,800 g for 30 min at 4°C, aliquoted the supernatants into four different tubes, and stored them at −80°C until assayed (<4 months). To get levels of antioxidant enzymatic activities in relation to proteins content in each sample, we determined total protein content using the autoanalyzer Cobas Integra 400 (Roche Diagnostics) with reagents purchased from RANDOX laboratory (Antrim, UK). We quantified the activity of glutathione reductase (in mU/mg protein), glutathione peroxidase (in mU/mg protein), superoxide dismutase (U/mg protein), and reduced-to-oxidised glutathione ratio using the autoanalyzer Cobas Integra 400 (Roche Diagnostics) with reagents purchased from RANDOX laboratory (Antrim, UK). Catalase activity (in U/mg protein) and malondialdehyde concentration (nmol/mL) were quantified according to standard colorimetric procedures (Cohen & Somerson, 1969; Galván et al., 2010; see also: Burraco et al., 2022b). Each sample was run in duplicate and intra-sample CV% were 3.44, 6.27, 8.77, 0.47, and 3.75 for catalase, glutathione reductase, glutathione peroxidase, superoxide dismutase, and malondialdehyde, respectively.

#### 2.4.3 Relative telomere length

We extracted liver and gut genomic DNA using the PureLink DNA extraction kit (Invitrogen, ThermoFisher) following the protocol provided by the manufacturer. We quantified the concentration and quality of DNA using a Nanodrop spectrophotometer. DNA was stored at −80 °C until assayed (<6 months).

Like in almost all vertebrates, *Xenopus laevis* telomere sequences consist of TTAGGG tandem repeats. To quantify variation in telomere length, we followed a procedure used in many taxa that provides relative telomere length measurements, based on the qPCR threshold cycle value of telomere repeats and a control single-copy gene. The genome of *X. laevis* presents almost no interstitial telomeric sequences (Meyne et al., 1990; Nanda et al., 2009), making the species a good candidate to estimate variation in telomere length through qPCR. We used primers designed for vertebrates to amplify their telomeric sequences: tel1b 5’-CGGTTTGTTTGGGTTTGGGTTTGGGTTTGGGTTTGGGTT-3’ and Tel2b 5’-GGCTTGCCTTACCCTTACCCTTACCCTTACCCTTACCCT-3’. We amplified a non-variable copy reference gene (RAG-1) as a control gene, using primers designed for *X. laevis* (Burraco et al., 2023a): XENRAG1-forward 5’-GCTCCATCGTCAGAGTTTAG-3’ and XENRAG1-reverse 5’-TGCTTCTGGTGGAGAATTAC-3’. For each gene, amplifications were conducted in a LightCycler 480 Real-Time PCR System (Roche) using a total of two 386-well plates including samples randomly distributed. Conditions for telomere qPCRs were 15 min at 95 °C for 15 s followed by 27 cycles of 95 °C for 15s, 58 °C for 30s and 72 °C for 30s, and a dissociation melting curve. Conditions for RAG qPCRs were 15 min at 95 °C followed by 40 cycles of 95 °C for 15s and 60 °C for 1 min, and a dissociation melting curve. Both primer combinations were used at 500 nM. Each qPCR plate contained a standard curve with five points serially diluted 1:1, a *no-template control* containing all reagents except DNA, and samples run in triplicate. Within each triplicate, we discarded samples with a cycle threshold values higher than 0.5. Following MIQE guidelines (Bustin et al., 2009). We also discarded a possible influence of DNA concentration or quality on relative telomere length measurements by running linear correlations between Nanodrop concentration (*R*^2^ = 0.001), 260/280 ratio (*R*^2^ = 0.009), and 260/230 ratio (*R*^2^ = 0.069). Intra-plate CV% was 0.71 and 0.43 for telomere and RAG amplifications, respectively.

### 2.5 Statistical analyses

All statistical analyses were conducted in R (R Development Core Team, version 4.21). We checked for data normality and homoscedasticity through Kolmogorov-Smirnov (*lillie.test* function*, nortest* package) and Breusch-Pagan tests (*bptest* function, *lmtest* package), respectively. To check for an effect of the different treatments on larval size, we conducted two linear models, one including *body mass*, and another including snout-to-vent length (SVL) as dependent variables. In both models, we included *fibre* and *stress* factors and their interaction as independent variables. We also explored the impact of fibre and stress exposure on gut mass by running a linear model with gut mass as the dependent variable, *fibre* and *stress* factors and their interaction as the independent variables, and *body mass* as the covariate. We then explored the possible effect of the experimental factors on the physiology of larvae by running linear models with *relative corticosterone concentration* (*i.e.,* corrected for body mass), oxidative stress parameters *(calatase, glutathione reductase, superoxide dismutase, glutathione peroxidase, lipid peroxidation*) or *relative telomere length* as dependent variables, and *fibre* and *stress* factors and their interaction as independent variables. When we detected a significant effect of either a single factor or the interaction between both factors, we conducted *post hoc* Tukey tests to identify differences between levels of the different factors using the function *emmeans* (package *emmeans*). To meet parametric assumptions, we log-transformed gut mass, relative corticosterone concentration, levels of oxidative stress parameters and relative telomere length.

## 3. Results

Fibre (*F*_2,91_ = 3.34, *P* = 0.040) and stress exposure (*F* _1,91_ = 7.03, *P* = 0.009) induced a reduction in larval body mass (Figure 1), whereas their interaction was not significant (*F* _2,91_ = 0.07, *P* = 0.931; Figure 1). MPs reduced larval body mass by 17.98% on average, as compared to control conditions (Tukey test = 0.034; Figure 1), whereas cellulose fibre did not affect larval body mass (Tukey test = 0.273; Figure 1). Likewise, CORT exposure reduced larval body mass by 15.49%, on average (Figure 1). SVL was also reduced by CORT (F_1,96_ = 7.76, *P* = 0.006), whereas fibre exposure or the interaction between both factors did not affect SVL (both *P* > 0.084). Gut mass, controlled for overall body mass, was reduced in response to CORT exposure by 25.33% on average (F_2,90_ = 11.01, *P* = 0.001), and it was unaffected by fibre exposure or the interaction between both factors (both *P* > 0.245).

**Figure 1.**
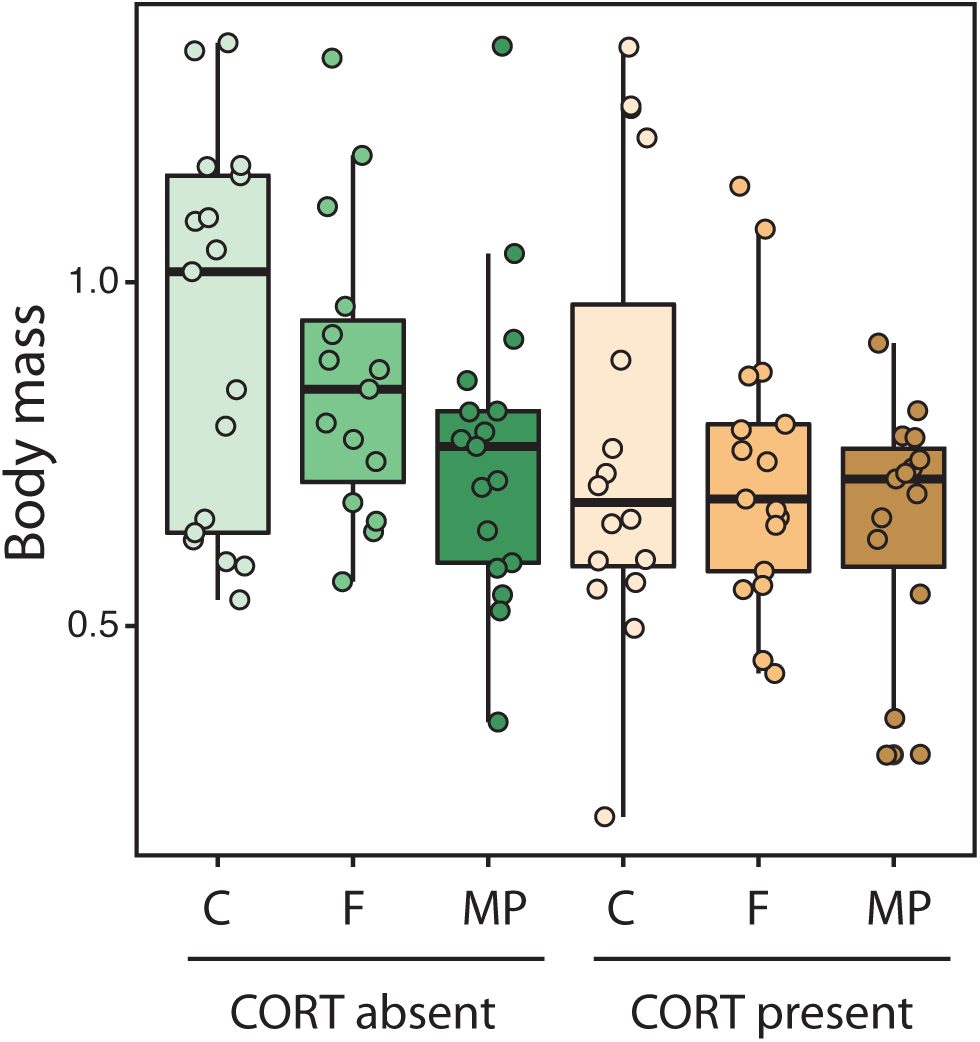
Variation in body mass in *Xenopus laevis* larvae exposed either to the presence or absence of exogenous corticosterone (100 nM CORT), combined with three different levels of fibre exposure (control with no fibres –C–, cellulose fibre –F–, or microplastic –MP– treatments, see Methods for details). Boxes represent 25^th^ to 75^th^ percentiles, lines within boxes represent median values, and vertical lines represent maximum and minimum data values.

Expectedly, larvae exposed to exogenous CORT had higher levels of the hormone in tail tissue (by 7.40% on average; F_1,82_ = 12.40, *P* < 0.001; Figure 2), but fibre exposure or the interaction between fibre and stress factors did not result in altered CORT levels (F_2,90_ = 0.52, *P* = 0.598 and F_2,90_ = 1.33, *P* = 0.268, respectively; Figure 2). Likewise, exogenous CORT increased the activity of the antioxidant enzymes catalase (F_2,90_ = 7.30, *P* = 0.008; Figure 3) and glutathione peroxidase (F_2,90_ = 12.37, *P* < 0.001; Figure 3), and the levels of lipid peroxidation (F_2,90_ = 18.07, *P* < 0.001; Figure 3). Fibre exposure or the interaction with exogenous CORT did not incur in redox alterations regarding the activities of antioxidant enzymes (all *P* > 0.737) or the markers of oxidative damage or status (malondialdehyde levels and the reduced-to-oxidised glutathione ratio, respectively; all *P* > 0.093). Finally, none of the experimental factors or their interaction caused significant reductions in telomere length, in either the liver or the gut (Figure 4; all P-values > 0.334).

**Figure 2.**
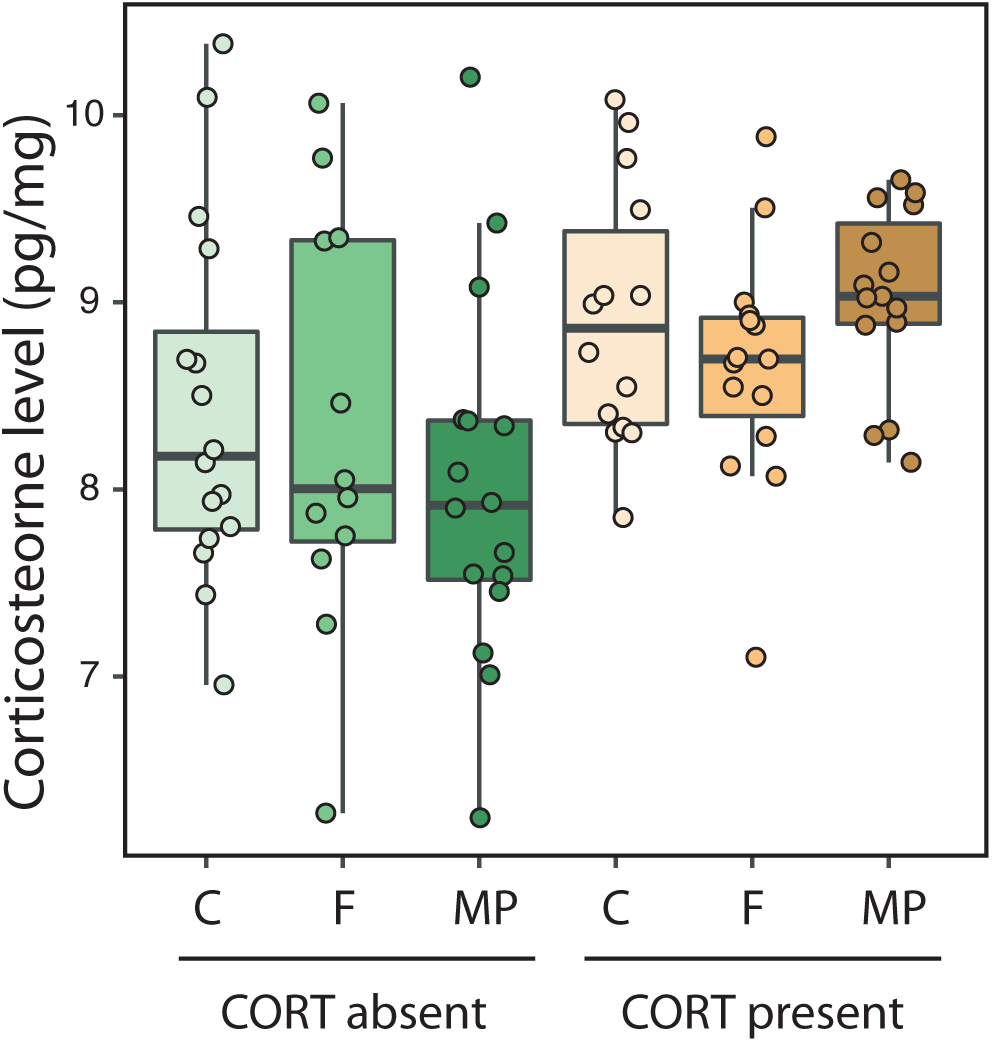
Variation in corticosterone concentration measured in tail tissue of *Xenopus laevis* larvae exposed either to the absent or present of corticosterone (CORT) exogenously added to the water (100 nM), combined with three different levels of fibre exposure (control with no fibres –C–, and cellulose fibre –F–, and microplastic –MP– treatments, see Methods for details). Boxes represent 25^th^ to 75^th^ percentiles, lines within boxes represent median values, and vertical lines represent maximum and minimum data values.

**Figure 3.**
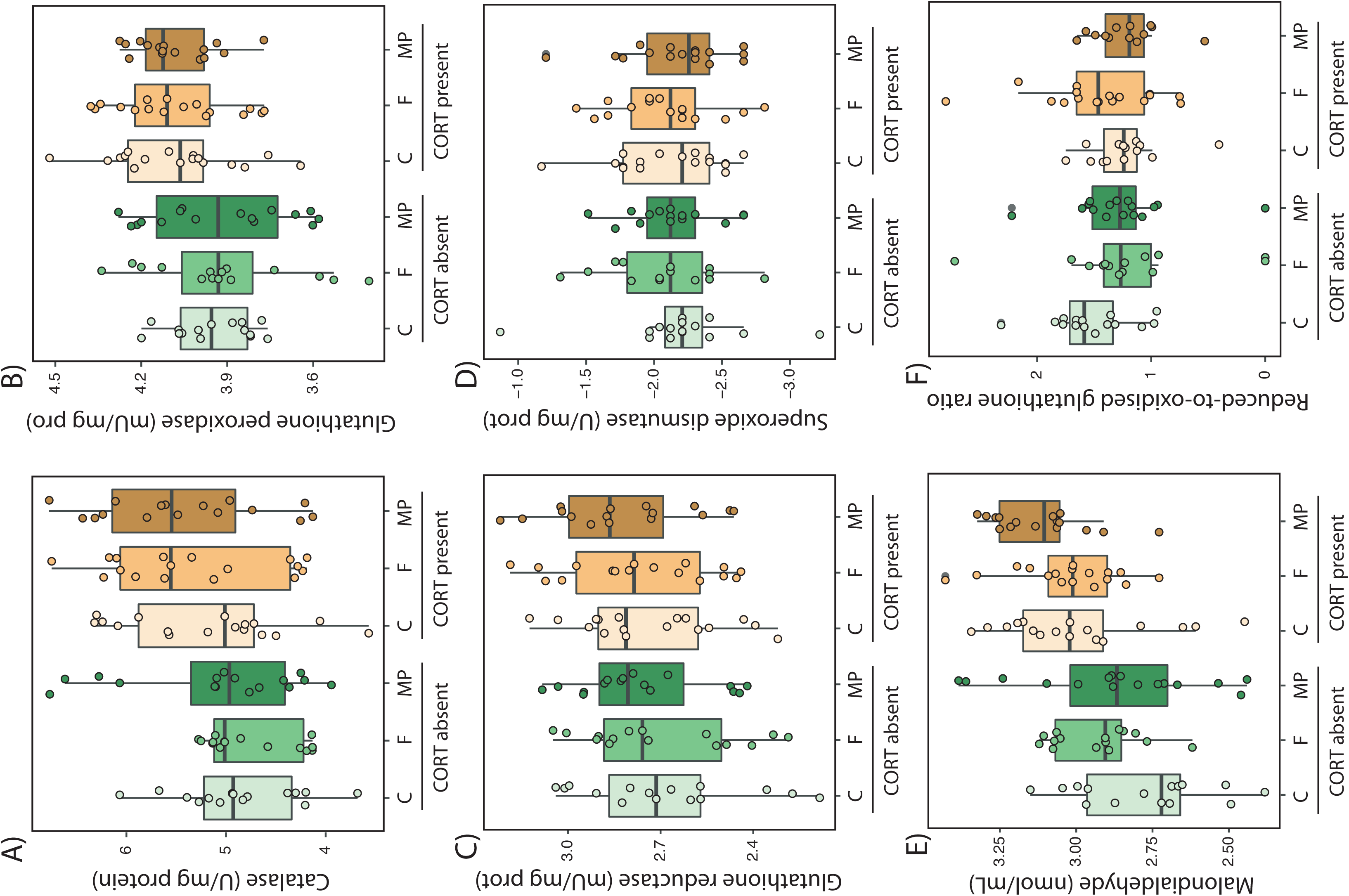
Variation in oxidative stress parameters. Antioxidant enzymes: A) catalase, B) glutathione peroxidase, C) glutathione reductase, and D) superoxide dismutase. Oxidative damage in lipids: E) malondialdehyde content. Non-enzymatic antioxidant status: F) reduced-to-oxidised glutathione). All these paremeters were measured in the liver of *Xenopus laevis* larvae exposed either to the absent or present of corticosterone (CORT) exogenously added to the water (100 nM), combined with three different levels of fibre exposure (control with no fibres –C–, and cellulose fibre –F–, and microplastic –MP– treatments, see Methods for details). Boxes represent 25^th^ to 75^th^ percentiles, lines within boxes represent median values, and vertical lines represent maximum and minimum data values.

**Figure 4.**
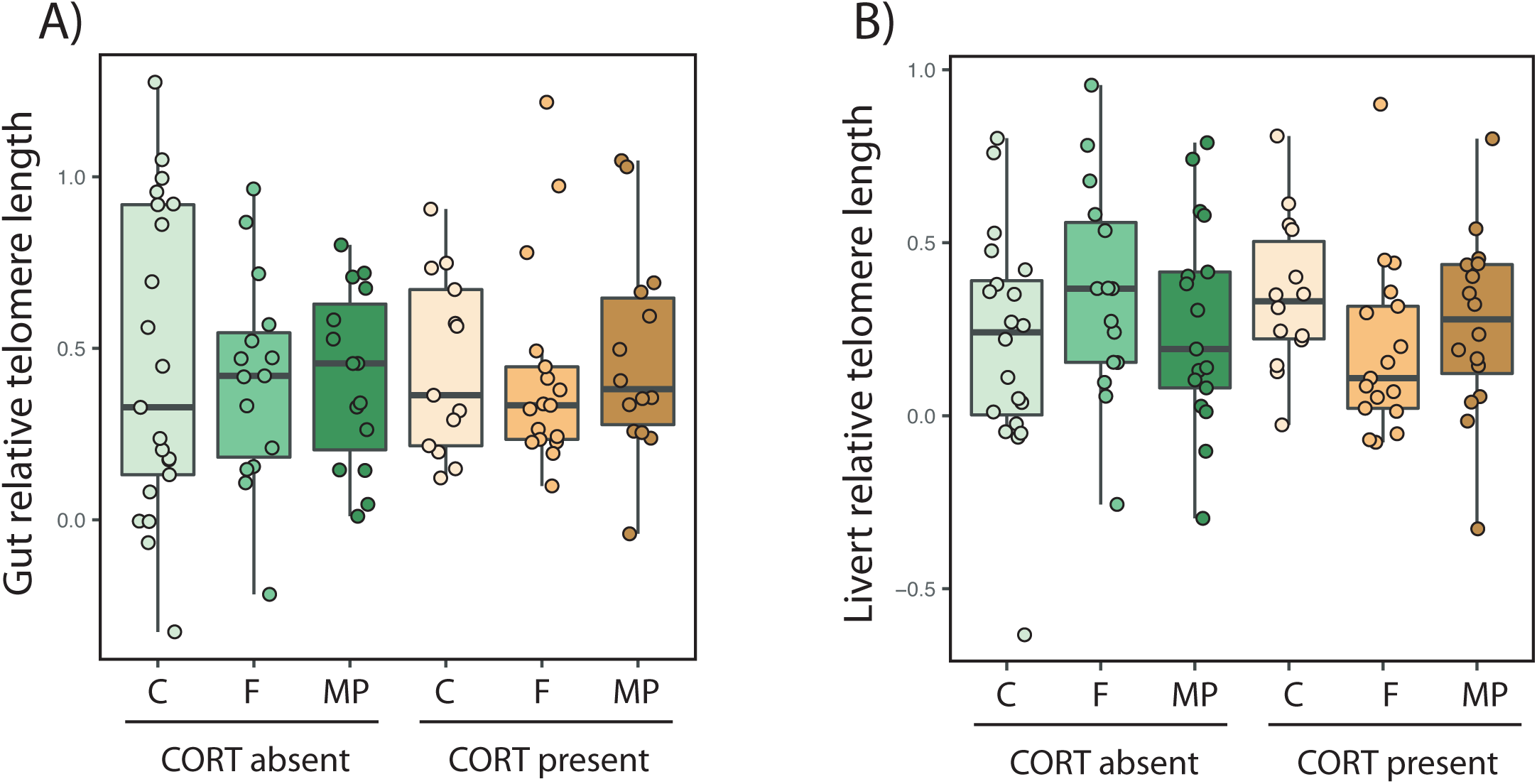
Variation in relative telomere length in the A) gut and B) liver of *Xenopus laevis* larvae exposed either to the absent or present of corticosterone (CORT) exogenously added to the water (100 nM), combined with three different levels of fibre exposure (control with no fibres –C–, and cellulose fibre –F–, and microplastic –MP– treatments, see Methods for details). Boxes represent 25^th^ to 75^th^ percentiles, lines within boxes represent median values, and vertical lines represent maximum and minimum data values.

## 4. Discussion

Microplastics (MPs) are expected to cause detrimental effects on animal development and health, however, the severity of the effect will vary across taxa and MPs load, and the mechanisms by which MPs can be harmful are not yet fully understood. Here we found evidence that MP exposure substantially reduced growth rate of the filter-feeding larvae of *Xenopus laevis*. However, despite their impact on growth rate, we did not observe major changes in physiological parameters associated with stress status (*i.e.,* corticosterone levels), or health and ageing (*i.e.,* redox balance and telomere length). Our findings, therefore, suggest that environmentally realistic concentration of MPs may have small impact on stress physiology even though they result in considerable growth related carry-over effects in amphibian larvae (but see: Ruthsatz et al., 2023b).

Coping with pollutants is often energetically demanding and can disrupt homeostasis in organisms inhabiting terrestrial or aquatic habitats (Noyes et al., 2009; rev. in Rohr et al., 2011). While eco-toxicological research has identified the causes and consequences of being exposed to some pollutants (García-Fernández et al., 2020; Michelangeli et al., 2022; Saaristo et al., 2018), further exploration is still needed in many other cases such as MP pollution. MPs were first identified in sea water in the 1960s (Bergmann et al., 2015; Carpenter & Smith, 1972) and, from that moment onwards, plastic litter has been found across marine, terrestrial, and atmospheric environments (Jiang, 2021; Prokić et al., 2019). Although MP exposure is expected to impair animal growth, development, or fecundity, evidence is equivocal across taxa (Chang et al., 2022; Prata et al., 2021; Zolotova et al., 2022). Our study shows a negative impact of MP exposure on the body mass of the filter-feeding larvae of *Xenopus laevis,* in accordance with other studies on aquatic organisms (rev. in Burgos-Aceves et al., 2022). Similar to other ectotherms, body mass is considered a good proxy for survival in amphibians, particularly at metamorphic stages, with smaller individuals often experiencing lower survival odds later in life (Berven, 1990; Cabrera-Guzmán et al., 2013; Gomez-Mestre & Tejedo, 2003; Smith-Gill & Berven, 1979). Intriguingly, we observed a lack of variation in skeletal growth (*i.e.,* body length) in larvae exposed to MPs. It is plausible that higher (but unrealistic) MP concentrations are needed to cause reductions in skeletal growth, and the interplay between MPs and other conditions such as temperature may also play a role here (Carreira et al., 2016). Additionally, it is possible that a plastic response in intestinal morphology such as increased gut length might have allowed for a compensation in skeletal growth as this is a mechanism often found in response to low food quality intake across taxa including amphibians (Ruthsatz et al., 2019b). If MPs are ingested together with the actual food source, the protein and energy density of the diet is reduced as MP particles are truly non-digestible fibres and lack any nutrients or energy that could be assimilated. In a previous study, we demonstrated that a plastic response towards a longer intestinal tract allowed for such a growth compensation in body mass but not body length when *Xenopus* larvae were exposed to a lower food quality through the ingestion of cellulose or MP fibres (Ruthsatz et al., 2022). However, MP exposure did not alter the mass of larval gut in the present study, suggesting that the response to MPs in terms of gut plasticity could be genotype or context dependent.

Once confirming the negative effects of MPs on the growth of *Xenopus laevis* larvae, we explored whether this process impaired their stress physiology. Chronically stressed animals can either upregulate or downregulate glucocorticoid levels (Rich & Romero, 2005; Romero & Beattie, 2022; Wingfield, 2013). In our study, larvae exposed to MPs for two weeks, did not experience changes in their CORT levels. This result could indicate that the MP concentration used in our study might not have been enough to induce metabolic alterations and to activate the hypothalamic-pituitary-interrenal axis and the subsequent glucocorticoid release in *Xenopus* larvae. Alternatively, larvae could have experienced unbalanced CORT levels soon after the exposure to MPs, but managed to recover basal hormone levels shortly afterwards. Tracking glucocorticoid dynamics in animals coping with MPs (and other pollutants) will help understand the actual impact of this relatively novel contaminant on the stress status of wildlife.

As expected, we observed that simulated stress through exogenously added CORT resulted in oxidative damage and elevated antioxidant responses in *Xenopus* larvae. In contrast, plastic pollution did not induce such changes in the redox status of larvae, not even in an interactive manner with previously physiologically stressed individuals. Based on meta-analytical evidence, pollution is known to lead to antioxidant responses or oxidative damage in several taxa, including amphibians (Chatelain et al., 2020; Isaksson, 2010; Martin et al., 2023). This also seems to be the case for MPs, however, oxidative stress changes may only occur in the short term or be dependent on MP concentration (Li et al., 2022; Li et al., 2023). Unlike the redox status, telomere dynamics have been overlooked in MP studies, which is surprising regarding the vast use of this ageing marker in eco-toxicological research (Chatelain et al., 2020; Salmón & Burraco, 2022). In amphibians, knowledge on telomere length variation in organisms coping with pollutants is, so far, restricted to a few studies (e.g. Cheron et al., 2022; Sabol et al., 2024; Zamora-Camacho et al., 2023), often observing little to none changes in telomere length. Likewise, in our experiment, MP exposure did not result in shortened telomeres, also in accordance with the observed lack of variation in CORT levels or oxidative stress. This result suggests that environmentally relevant concentration of MPs do not alter accelerate ageing rates in *Xenopus laevis* larvae. Alternatively, reductions in larval growth might have resulted in reduced cell division and less demanding metabolic processes required for somatic maintenance, which could have finally resulted in unaltered telomeres. This possibility should be explored through estimates of cell division rates and aerobic (mitochondrial) metabolism across tissues.

## 5. Conclusions

Our study confirms that the exposure to an environmentally realistic concentration of MPs reduces growth in filter-feeding amphibian larvae. Such growth effects do not include consequences for stress and ageing-related physiological mechanisms, suggesting minor carry-over effects of MP exposure. Overall, this study improves our mechanistic understanding of the effect of MPs on the life history and physiology of aquatic organisms, a knowledge that could be integrated into mechanistic niche modelling to finally develop effective conservation actions (e.g. Burraco et al., 2022a; Kearney & Porter, 2009; Newman et al., 2022).

## 6. Data accessibility

The dataset of this study can be accessed at https://doi.org/10.6084/m9.figshare.25958923.v1

## 7. Author contributions

**CM**: Data curation (supporting); Methodology (equal); Investigation (equal); Writing – original draft (equal); Writing – review and editing (equal). **KR**: Conceptualization (lead); Supervision (equal); Methodology (equal); Data curation (equal); Formal analysis (supporting); Investigation (equal); Writing – original draft (equal); Writing – review and editing (equal); Project administration (equal); Funding acquisition (lead). **IGM**: Conceptualization (equal); Resources (supporting); Writing – review and editing (equal); analysis (lead); Investigation (equal); Methodology (equal); Project administration (equal); Writing – original draft (equal); Writing – review and editing (equal); Funding acquisition (supporting).

## 8. Ethics statement

All experimental procedures were conducted at Doñana Biological Station (Spanish National Research Council, CSIC, Seville, Spain) following the approved bioethical permit 12/12/2023/108.

## 11. Acknowledgements

We are grateful to M. Gutierrez, F. Miranda, and Clara for assistance in the Ecophysiology laboratory at the Doñana Biological Station. We thank B. Oetken for his help preparing the experiments in Braunschweig.

